# Species phylogenetics using intraprotein residue dependence inferred with protein foundation models

**DOI:** 10.64898/2026.05.26.727927

**Authors:** Rohan Alibutud, Mohammad Erfan Mowlaei, Sudhir Kumar

## Abstract

Traditional molecular phylogenetic inference uses multiple sequence alignments and amino acid substitution models that generally assume that sites evolve independently. In contrast, protein foundation models (pFMs) are deep learning models that reveal residue dependence patterns within sequences. There are two types of pFMs: Evolutionary Scale Models (esm-pFM) for analyzing individual protein sequences, and MSA Transformer (msat-pFM) models for analyzing MSAs. Using pFMs to analyze sequences produces residue attention matrices (*RAM*s) and sequence attention matrices (*SAM*s), respectively. *RAM*s and *SAM*s can be used to estimate pairwise distances and, thus, reconstruct phylogenies using the Neighbor-Joining (NJ) method. Here, we compare the performance of *RAM*-NJ and *SAM*-NJ phylogenies in reconstructing established species relationships across different proteins and taxon sets. Surprisingly, *RAM*-NJ phylogenies were generally as accurate as *SAM*-NJ phylogenies even though *RAM* inference does not require construction or analysis of an MSA. Our results show that pFM-based analyses can recover species relationships without conventional substitution models or, in the case of *RAM*s, a multiple sequence alignment. These results suggest that evolutionary relationships are reflected not only in primary sequence differences between species, but also in the sequence-specific patterns of inter-residue dependence produced by pFM analysis of individual sequences. However, pFM phylogenies are less accurate than those obtained with standard maximum likelihood approaches, underscoring the need to further develop methods to harness *RAM*s and *SAM*s more effectively.

## Introduction

Molecular phylogenetic inference traditionally begins with a multiple sequence alignment (MSA) in which amino acid residues in homologous positions are compared across species. Researchers then model their evolution using stochastic models that describe amino acid frequencies and substitution rates, as well as variation in site-specific substitution rates (Nei and Kumar 2000; Yang 2006). Maximum likelihood and distance methods use substitution models that generally assume that sites evolve independently and that amino acid substitution processes are stationary, reversible, and time-homogeneous (Kumar et al. 2026). These assumptions make statistical analysis computationally tractable but do not capture the context dependence of amino acid substitutions, which arises because the fitness and evolutionary fate of a substitution depend on residues at other positions in the protein (Kumar et al. 2026; Schmelkin et al. 2026).

Protein foundation models (pFMs), also called protein language models, offer an alternative approach that focuses on residue dependence across sites within a protein rather than variation among sequences (Kumar et al. 2026). Researchers estimate (train) parameters (weights) of these models on millions of naturally occurring proteins using transformer architectures with many layers and attention heads (Devlin et al. 2019; Rao et al. 2021; Rives et al. 2021). During training, they identify statistical regularities in protein sequence corpora, including patterns associated with sequence composition, structural constraints, functional motifs, residue context, and homology (Devlin et al. 2019; Rao et al. 2021; Rives et al. 2021). Pre-trained pFMs have subsequently been used for protein structure prediction, functional annotation, variant-effect prediction, protein design, and other biological tasks (Sapoval et al. 2022; Schmirler et al. 2024; Braichenko et al. 2025).

In this study, we examine two major pFMs, which differ in how they process sequence information during training and analysis. One is MSA Transformer, which is trained on a large collection of MSAs (Rao et al. 2021) that we call msat-pFM (Kumar et al. 2026). When an MSA is analyzed with a pre-trained msat-pFM, the model produces sequence attention matrices (*SAM*s) describing relationships among sequences at homologous alignment positions. Because msat-pFM directly analyzes aligned homologs, it might be expected to extract signals of evolutionary relationships among sequences. Indeed, *SAM* elements have been shown to correlate with pairwise sequence divergence and, after supervised fine-tuning, to predict Hamming distance between sequences (Lupo et al. 2022). A simple transformation of *SAM*s, without fine-tuning, can construct a phylogeny using the neighbor-joining (NJ) method (Saitou and Nei 1987), resembling phylogenies inferred from conventional distances (Chen et al. 2026). Therefore, *SAM*s contain significant phylogenetic information. The second is trained by using individual sequences one by one during Evolutionary Scale Modeling (Rives et al. 2021), with the resulting model called esm-pFMs (Kumar et al. 2026). During training, the model receives no explicit information about sequence homologies or MSAs, species identities, or protein functions. When an individual protein sequence is analyzed with a pretrained esm-pFM, the model produces residue attention matrices (*RAM*s), whose elements quantify context dependence among residues within the same sequence (Kumar et al. 2026). The use of pFM outputs from individually analyzed sequences to investigate evolutionary relationships has begun, including studies based on sequence embeddings and other pFM-derived quantities (Tule et al. 2025; Stakauskas and Górecki 2026). However, the potential of *RAM*s themselves for reconstructing species relationships has not been systematically evaluated.

However, using these pFMs to infer species relationships of higher organisms remains completely underexplored, and several important questions remain unresolved. First, previous studies have focused on relationships among members of protein families, including paralogs (Tule et al. 2025; Stakauskas and Górecki 2026), for which no ground truth exists, unlike the recovery of accepted species relationships using orthologous sequences. Second, researchers have not compared the performance of *SAM* phylogenies, which require an MSA, with that of *RAM* phylogenies, which do not. Third, little is known about how species-relationship information is distributed across the heads and layers that comprise the pFM neural network, beyond limited reports that *SAM*s produced by intermediate layers show a higher correlation with pairwise similarity among sequence homologs (Lupo et al. 2022).

Here, we investigate the phylogenetic signal from proteins analyzed using these two contrasting pre-trained pFMs. For msat-pFM, we estimate evolutionary distance between sequences directly from *SAM*s generated from an MSA, using a modified version of the method described by Chen et al. (Chen et al. 2026). To build phylogenies using *RAM*s derived from individual sequence analyses, we introduce an approach that first aligns each sequence pair to establish residue-level evolutionary correspondence such that *RAM*s can be used to estimate pairwise distances (see *Results*). We do this independently for each sequence pair to build an attention distance matrix between sequences. We use *SAM* and *RAM* distance matrices to generate NJ phylogenies (see *Results*).

We evaluate and compare the accuracy of NJ phylogenies from *RAM* and *SAM* distances for different proteins, using datasets from four closely and distantly related species and a larger collection of mammalian species. We evaluate accuracy by comparing them with accepted species relationships and with phylogenies inferred using a conventional maximum likelihood (ML) approach. We also examine phylogenetic informativity of *RAM*s and *SAM*s across pFM heads and layers. Our results show that *RAM* phylogenies contain strong phylogenetic signals and are, unexpectedly, as accurate as *SAM* phylogenies, despite the former not relying on multiple sequence alignment. We suggest that *RAM*s produced by esm-pFM for individual sequences retain imprints of species evolutionary relationships.

## Results

### Estimating evolutionary distances from *RAM*s and *SAM*s

We introduce two methods for reconstructing phylogenetic trees from the attention matrices of esm-pFM and msat-pFM. For esm-pFM, we produce *RAM*s for each attention head in each model layer when analyzing individual sequences. Each element of a *RAM* describes the relative attention from one residue position to another. Consider two orthologous sequences, A and B, of lengths *a* and *b* (**Fig. 1a**). For attention head *h*, esm-pFM produces *RAM*s (*R*_A_ and *R*_B_) of dimensions *a*×*a* and *b*×*b*, respectively (**Fig. 1a**).

**Figure 1.**
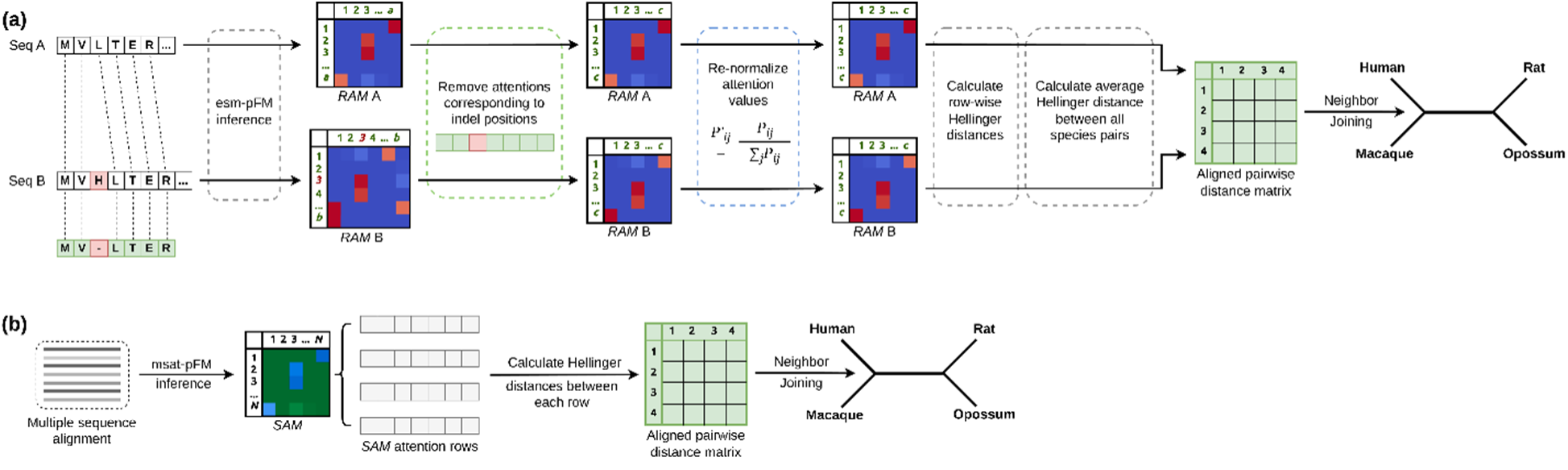
Estimation of phylogenetic distances from protein foundation model attention matrices. (**a**) Analysis of an individual protein sequence with a pre-trained esm-pFM produces a residue attention matrix (*RAM*) for each attention head and layer. Orthologous sequences are analyzed independently, and pairwise sequence alignments are then used to identify homologous residues between sequences A and B. *RAM* elements involving positions aligned to an insertion or deletion are removed, which is followed by calculating row-wise Hellinger distances to estimate *RAM* distance as shown in equations 1-3. Distances calculated between all sequence pairs are assembled into a matrix from which a neighbor-joining (NJ) phylogeny is inferred. (**b**) Analysis using the pre-trained msat-pFM begins with a multiple sequence alignment (MSA). Sequence attention matrices (*SAM*s) are extracted, and then individual rows are compared against each other using row-wise Hellinger distances. Each row corresponds to a different species sequence, producing a pairwise distance matrix used to infer NJ phylogenies.

*RAM*s cannot be compared directly because residue homology cannot be assumed from position alone when comparing individual sequences. Thus, we first align sequences A and B, e.g., by using the Needleman-Wunsch (1970) algorithm. We align each sequence pair once and use the residue alignment when comparing *RAM*s. Positions aligned to gaps (insertions/deletions) in either sequence are removed from the corresponding rows and columns of both *R*_A_ and *R*_B_. This produces two reduced *R*_A_ and *R*_B_ matrices of dimension *c*×*c*, where *c* is the number of alignment positions containing valid amino acids in both sequences. Thus, each retained matrix element describes attention between pairs of positionally homologous residues (**Fig. 1a**).

Each row of a *RAM* shows how a residue at a given site attends to every other site, with attention weights that sum to 1, analogous to probabilities. Pruning matrix elements that correspond to gaps in a *RAM* also removes the corresponding weights, so we add a renormalization step to ensure the remaining attention weights sum to 1. For sequence A, the renormalized attention from retained residue *i* to retained residue *j* is:

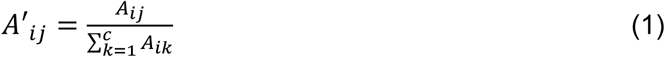

We then calculate the distance between the two matrices using the Hellinger distance (*H*) (Hellinger 1909), which measures differences in corresponding attention probabilities that sum to 1 in a given row in *RAM*_A_ and *RAM*_B_, rather than Euclidean distance, which is better suited to unconstrained numerical vectors. Furthermore, Euclidean distances are dominated by high-probability elements in vectors (Tule et al. 2025), and cosine distances can yield negative values and thus require further transformations (Nayar et al. 2025).

Hellinger distance based on the attention probabilities in a given row (residue *i*) in *RAM*_A_ and *RAM*_B_ is given by the following equation:

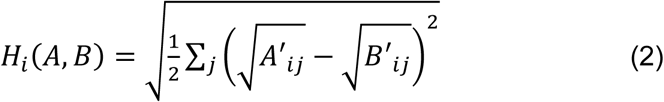

Here, *A’_ij_* denotes the normalized attention from residue *i* to residue *j* in *RAM*_A_, and *B’_ij_*denotes the normalized attention from residue *i* to residue *j* in *RAM*_B_. The Hellinger equation computes a scaled Euclidean distance between the square roots of *RAM*_A_ and *RAM*_B_ probabilities. Using the square root effectively compresses differences between large attention weights while increasing the relative contribution of differences among smaller weights.

The evolutionary distance based on all *c* residues is then given by the root-mean-square of Hellinger distances averaged over *c* rows:

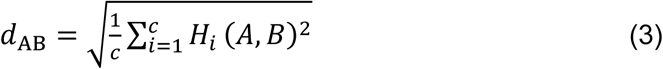

Thus, *d*_AB_ summarizes the divergence in residue interaction patterns across the homologous positions between two sequences.

Repeating this process for every pair in a set of sequences yields a pairwise distance matrix. For a pFM with *L* layers and *H* attention heads per layer, the procedure above produces *L*×*H* pairwise *RAM* distance matrices. We infer an NJ tree for each pairwise distance matrix using its lower triangle, since it is symmetric. For the esm-pFM checkpoint used here, this yielded 240 *RAM* phylogenies across 12 layers and 20 heads per layer.

We estimated *SAM* distances as a simple modification of the above approach (see *Materials and Methods*), producing 144 *SAM*s and 144 *SAM* phylogenies for each input MSA, corresponding to the 12 layers and 12 heads of the msat-pFM model. Because the *SAM*s also consist of post-softmax attention weights, row entries are analogous to probability distributions and are used to calculate Hellinger distances as above (see equations 2 and 3).

### Accuracies of pFM phylogenies for four taxa datasets

As a test of pFMs’ ability to reconstruct a simple phylogeny, we first examined transferrin orthologs from four distantly related mammals (**Fig. 2a**). These species have an unambiguously accepted relationship in which humans (*Homo sapiens*) and macaques (*Macaca fascicularis*) form a clade, the rat (*Rattus norvegicus*) is their more distant placental relative, and the opossum (*Monodelphis domestica*) is the marsupial outgroup. The retrieved transferrin sequences ranged from 628 amino acids in opossum to 695 in macaque. *RAM* phylogenies were correct for all 240 layer-heads (100% accuracy). In contrast, 6 of 144 *SAM* layer-head phylogenies were incorrect (95.8% accuracy).

**Figure 2.**
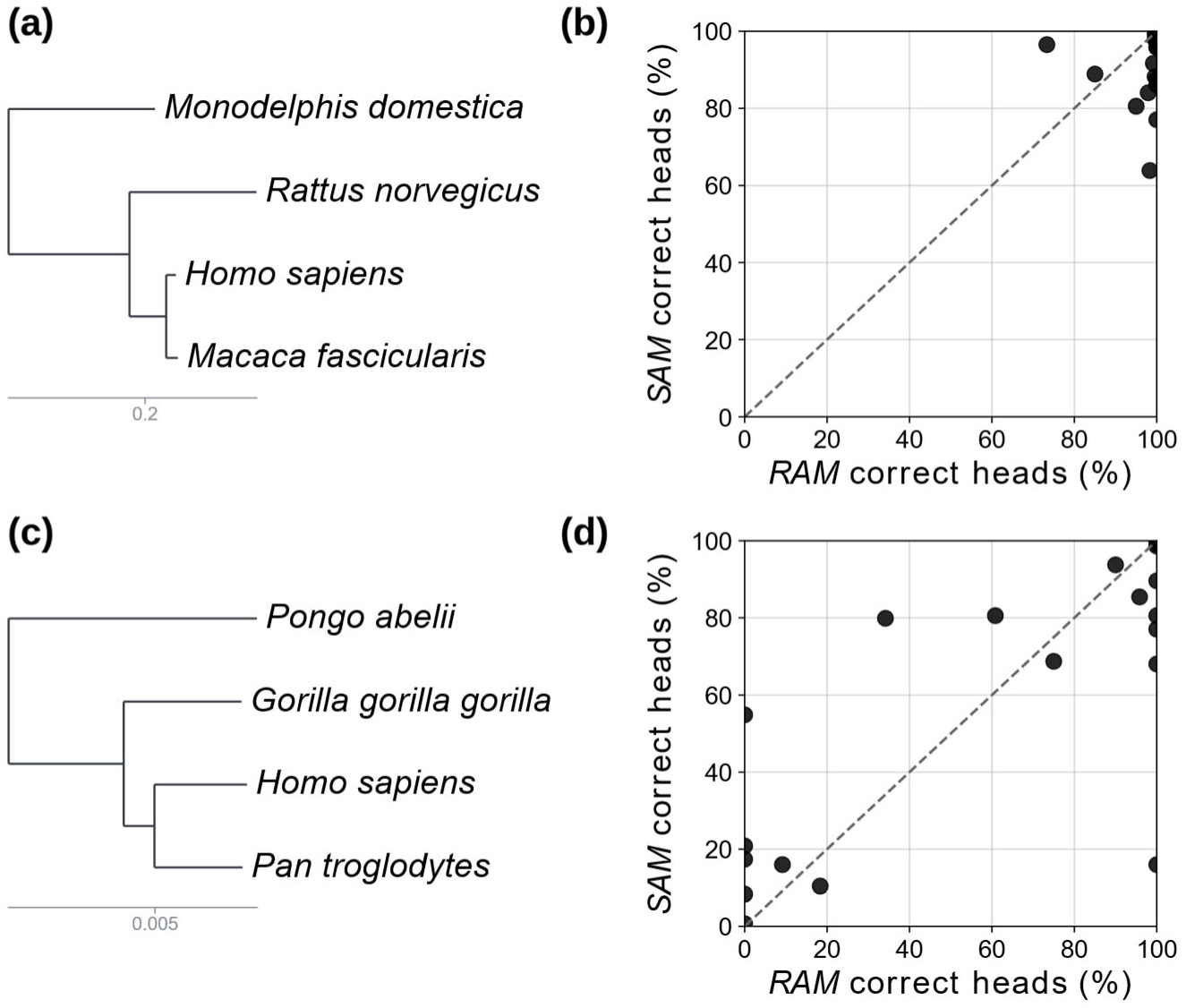
Fraction of attention heads recovering the two four-species phylogenies correctly. (**a**) Standard species relationship among humans (*Homo sapiens*), crab-eating macaques (*Macaca fascicularis*), rats (*Rattus norvegicus*), and gray short-tailed opossums (*Monodelphis domestica*). The phylogeny is drawn to scale based on the branch lengths available from the OrthoMaM resource. (**b**) Percentage of *RAM* and *SAM* layer-heads phylogenies producing the correct phylogeny in the panel for 20 orthologous proteins. **(c)** Standard species relationships among humans (*Homo sapiens*), chimpanzees (*Pan troglodytes*), gorillas (*Gorilla gorilla*), and orangutans (*Pongo abelii*). The phylogeny is drawn to scale based on the branch lengths available from the OrthoMaM resource. The short interval between the human-chimpanzee and gorilla clades makes this relationship harder to resolve from individual proteins and can also produce genuine gene-tree discordance. **(d)** Percentage of *RAM* and *SAM* head-specific phylogenies recovering the correct phylogeny in panel *c* for 20 orthologous proteins.

We tested whether these patterns were general by expanding the analysis to 20 orthologous proteins sampled from the OrthoMaM database (see *Materials and Methods*). For this 4-species inference, *RAM* layer-head phylogenies were more accurate than *SAM* phylogenies for 17 of 20 proteins (**Fig. 2b**). *RAM* layer-head phylogenies were correct 97.3% of the time, compared with 91.2% for *SAM*s.

The preceding analysis involved distantly related species with relatively long internal branches, making phylogeny comparatively easy to reconstruct. As a more difficult example, we chose the 4-taxon phylogeny of human, chimpanzee (*Pan troglodytes*), gorilla (*Gorilla gorilla*), and orangutan (*Pongo abelii*) (**Fig. 2c**). We expect lower accuracy in this case because of the short internodal distance separating the human-chimpanzee lineage from the gorilla lineage and the known potential for incomplete lineage sorting (ILS) in this part of the primate phylogeny (Blom et al. 2026). In this case, we expected MSA-based inference to be more powerful, but the fraction of correct *RAM* layer-head phylogenies (59.2%) was similar to *SAM* layer-head phylogenies (58.3%) (**Fig. 2d**). Interestingly, the fraction of correct *RAM* layer-head phylogenies was 0 for 5 of the 20 proteins (APOBEC2, COCH, GPX3, TF, UBA2), but none for *SAM*s. So, using an MSA seems helpful for some proteins.

### Phylogenetic accuracy in larger mammalian datasets

Phylogenetic analyses of four-taxon datasets select the true tree from only three possible unrooted topologies and, therefore, offer limited resolution for comparing phylogenetic methods. We reasoned that an explicit MSA might become more beneficial as the number and diversity of sequences increased. We therefore expanded each of the 20 protein datasets to 34 mammalian species (**Fig. 3a**). These taxa span a broad range of evolutionary timescales, from approximately 0.45 million years between closely related bovids to approximately 160 million years between placental mammals and the marsupial outgroup (Kumar et al. 2022). We used the OrthoMaM genome-wide phylogeny as the reference species tree (Allio et al. 2024). We quantified accuracy as the percentage of correct clades in the layer-head phylogeny (PCC; see *Methods*). *RAM*s and *SAM*s performed similarly on this dataset, with mean PCCs of 39.8% and 39.2%, respectively (**Fig. 3b**; paired t-test, p = 0.491).

**Figure 3.**
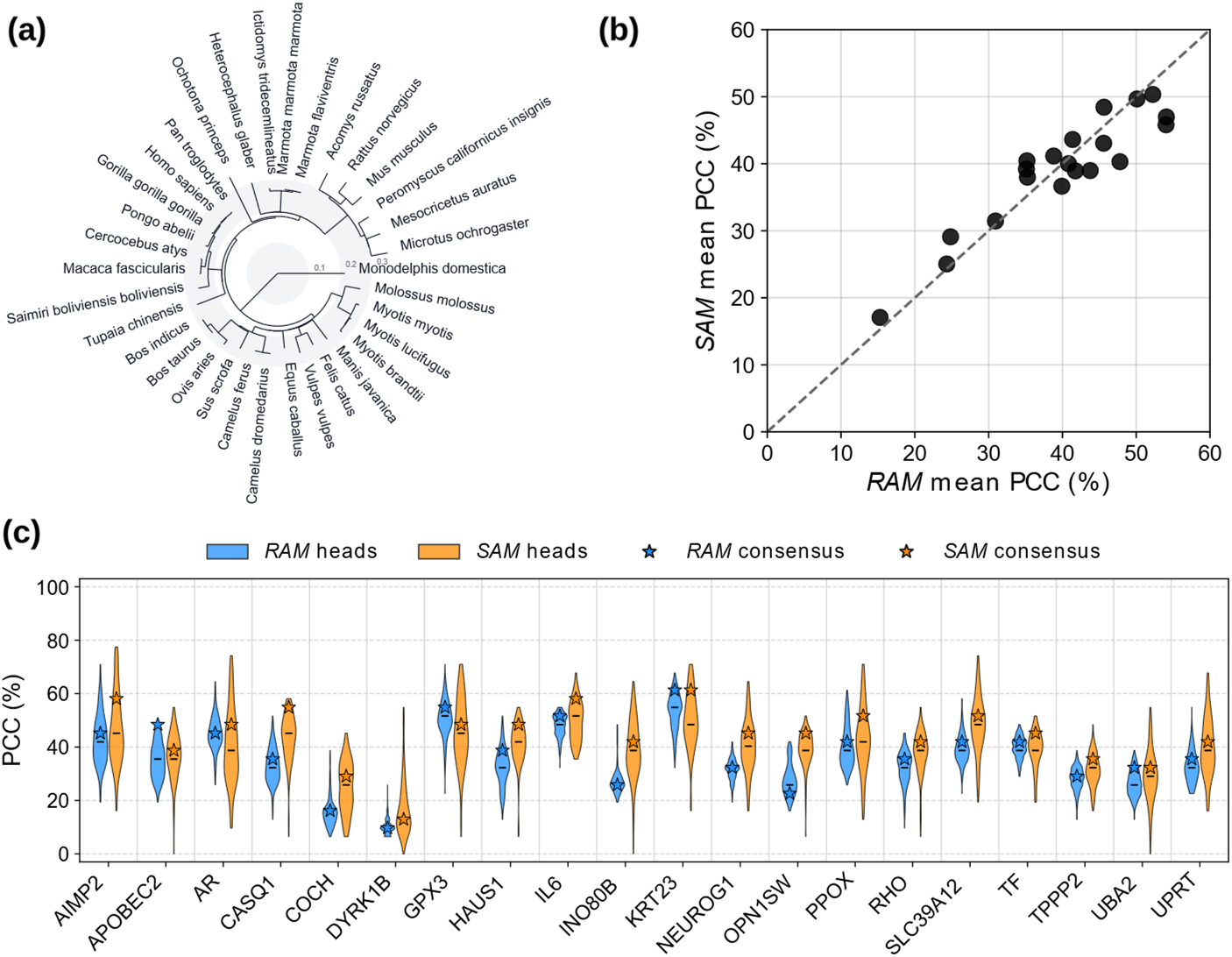
Accuracy of pFM-based phylogenetic inference for a larger tree. (**a**) Reference phylogeny for the 34 mammalian species analyzed, with topology obtained from OrthoMaM. (**b**) A scatterplot showing the percentage of clades correct (PCC) in *RAM* and *SAM* phylogenies for 20 orthologous proteins. (**c**) Violin plot showing the range in PCC of the *RAM* and *SAM* phylogenies across attention heads, with the stars denoting the PCC of the all-layers combined phylogeny for that protein.

### Phylogenies across layers and proteins

We next asked how well the *SAM* and *RAM* approaches reconstructed the 34-species phylogeny when information was combined across all 20 proteins. As an additional comparison, we inferred maximum likelihood (ML) trees for the same 20 proteins as a reference for classical phylogenetic methods (see *Materials and Methods*). For *RAM* and *SAM* analyses, we combined the 20 protein trees using ASTRAL to infer a final multi-protein phylogeny for each method. We compared each resulting tree with the OrthoMaM reference phylogeny using tanglegrams (**Fig. 4**). ASTRAL reported local posterior probabilities at internal branches, which indicate support among the 20 protein phylogenies for each inferred relationship. Thus, the topology indicates accuracy relative to the reference tree, whereas the posterior probability indicates how consistently that relationship is supported by the protein-specific trees.

**Figure 4.**
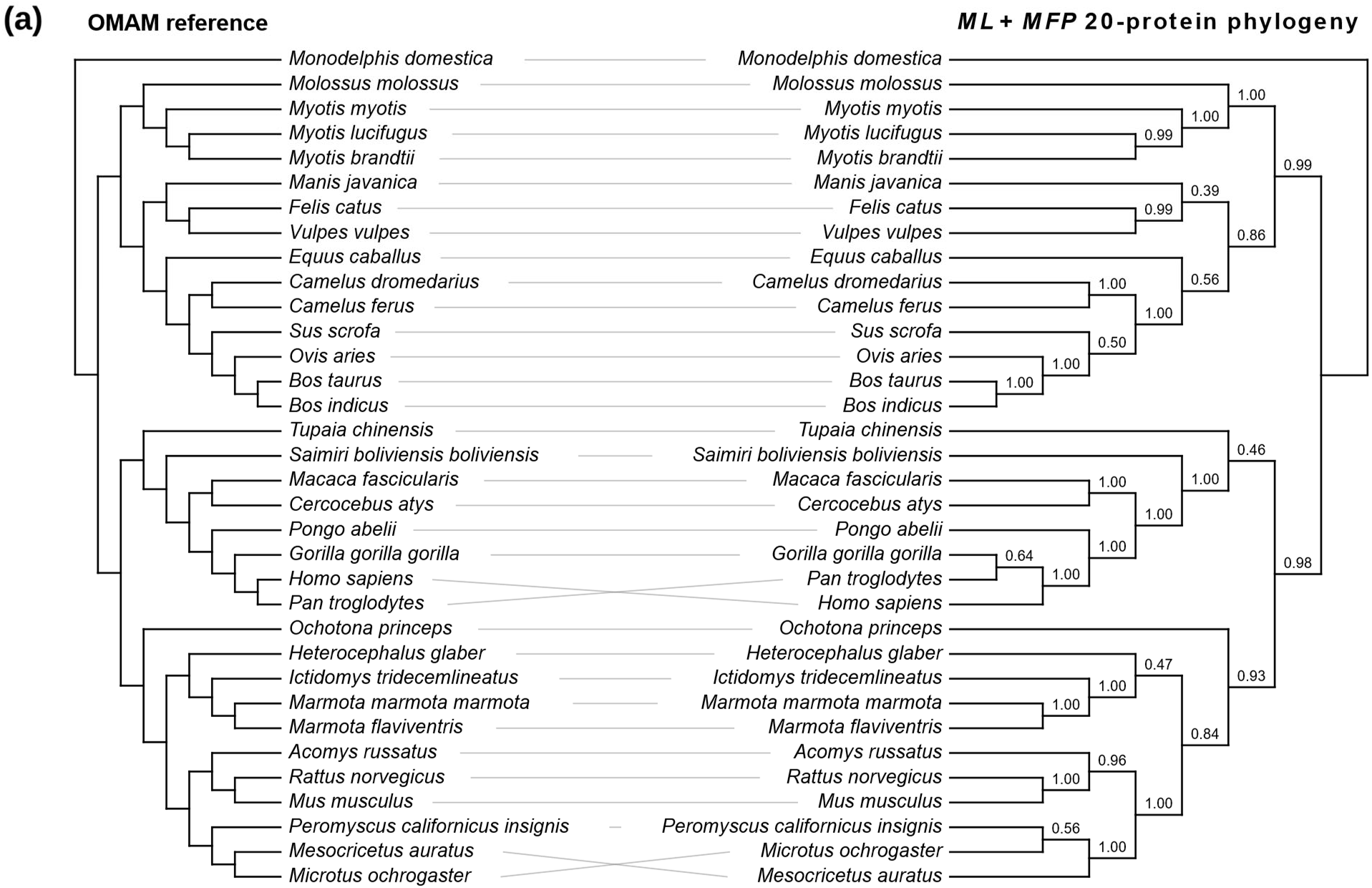
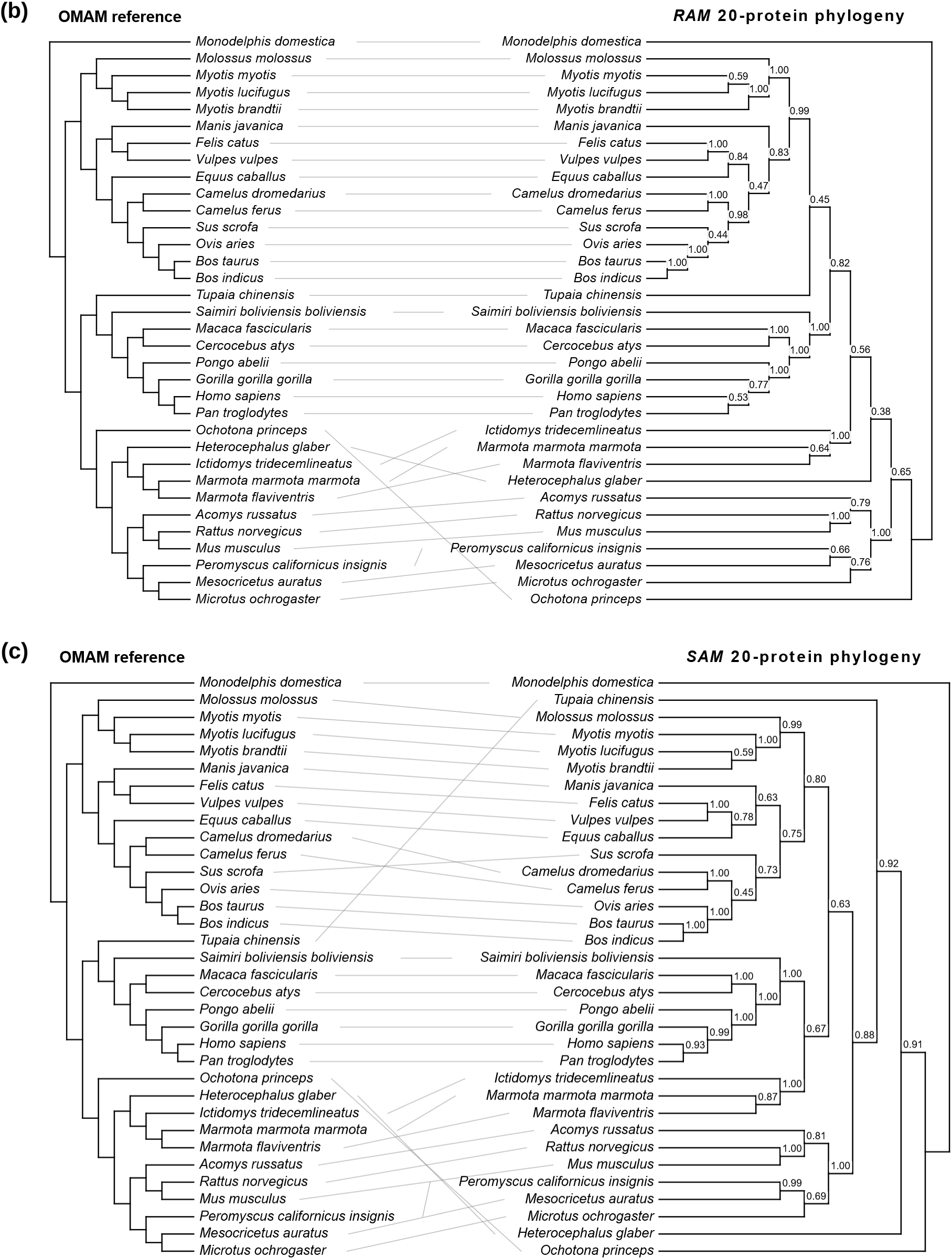
Comparison of 34-species 20-protein trees against the OrthoMaM reference phylogeny. The reference tree was obtained by pruning the OrthoMaM SuperMatrix tree to the 34 species used in the analysis. Crossovers indicate tips that are mismatched relative to the reference tree once both were rooted on the outgroup (*Monodelphis domestica*). **(a)** Tanglegram comparing the reference tree against the ML phylogeny. **(b)** Tanglegram comparing the reference tree against the 20-protein *RAM* phylogeny. **(c)** Tanglegram comparing the reference tree against the 20-protein *SAM* phylogeny. Local posterior probabilities from ASTRAL analysis are shown for each tree.

The ML consensus phylogeny showed the greatest overall agreement with the reference tree (**Fig. 4a**). Most major mammalian groups and many relationships within them were recovered with high posterior probability. For example, relationships among the sampled bats, bovids, and most primates had posterior probabilities near 1.0. However, ML did not recover every expected relationship. Most notably, it grouped *Pan troglodytes* with *Gorilla gorilla* rather than with *Homo sapiens*. Importantly, this incorrect relationship had only moderate posterior support (0.64), even though surrounding primate relationships were strongly supported. A similar pattern occurred among rodents. The reference phylogeny places *Mesocricetus auratus* and *Microtus ochrogaster* as sister taxa, whereas the ML consensus grouped *Microtus* with *Peromyscus californicus*. This incorrect grouping also received relatively weak support (0.56), whereas several neighboring rodent relationships received posterior probabilities greater than 0.95.

The *RAM* consensus phylogeny also recovered many of the major relationships in the reference tree despite being assembled from phylogenies inferred without an MSA or an explicit amino acid substitution model **(Fig. 4b)**. For example, the relationships among the three *Myotis* species, the *Felis*-*Vulpes* grouping, the two *Camelus* species, the bovids, the *Macaca*-*Cercocebus* pair, and the *Rattus*-*Mus* pair were recovered with high posterior probabilities, many approaching or reaching 1.0. *RAM* also correctly recovered the *Homo sapiens*-*Pan troglodytes* clade that was missed by the ML consensus. Support for this relationship, however, was modest (0.53), whereas the subsequent grouping with *Gorilla gorilla* had higher support (0.77), and the broader hominid grouping, including *Pongo abelii* was strongly supported (1.00). Thus, *RAM* recovered the accepted hominid topology, but the posterior probabilities indicate that the human–chimpanzee relationship was among the less consistently recovered relationships across the 20 proteins.

Most disagreements between the *RAM* consensus and the reference tree occurred within rodents, where several closely spaced relationships were difficult to reconstruct. *RAM* correctly recovered the *Rattus norvegicus-Mus musculus* grouping with a high posterior probability (1.00) and the broader grouping including *Acomys russatus* with a moderate support (0.79). However, it grouped *Peromyscus californicus* with *Mesocricetus auratus* rather than recovering the reference *Mesocricetus-Microtus* relationship. This discordant grouping had only moderate posterior support (0.66), as did the next deeper relationship (0.76). More generally, several deeper branches in the *RAM* consensus had only moderate posterior probabilities (0.4-0.8). Therefore, the *RAM* consensus recovered many well-established local clades with strong support, while uncertainty was concentrated primarily in deeper or more difficult parts of the phylogeny.

The *SAM* consensus phylogeny likewise recovered many major species relationships but showed more conspicuous departures from the reference topology **(Fig. 4c)**. Within primates, *SAM* correctly recovered the *Homo sapiens-Pan troglodytes* relationship with a posterior probability of 0.93, substantially higher than the corresponding *RAM* support of 0.53. The successive inclusion of *Gorilla gorilla* and *Pongo abelii* was also strongly supported (> 0.95), and the *Macaca-Cercocebus* relationship also received similarly high support. Thus, for these well-recovered primate relationships, *SAM* produced both the correct topology and strong support. However, the *SAM* consensus incorrectly placed *Tupaia chinensis* relative to the primate clade and differed from the reference tree in its placement of *Sus scrofa*. Several rodent relationships were also incorrect. In particular, *SAM* grouped *Peromyscus californicus* and *Mesocricetus auratus* with very high posterior probability (0.99), which conflicts with the reference tree and is a strongly supported false inference.

Comparing the three consensus trees reveals differences that topology alone does not capture. ML produced the most accurate overall consensus tree, and its principal incorrect relationships tended to have only moderate posterior support. *RAM* recovered many of the same established clades and correctly reconstructed the human-chimpanzee relationship missed by ML, although support for some deeper and more difficult *RAM* relationships was lower. *SAM* also recovered many local clades, including the human-chimpanzee relationship, but it also strongly supported several erroneous clade assignments. Thus, posterior probabilities help distinguish two different causes of disagreement with the reference tree: relationships that remain uncertain across proteins, as seen for several *RAM* and ML discrepancies, and relationships that are reproducibly inferred by a method but disagree with the accepted species topology, as illustrated by some *SAM* relationships.

### Distribution of phylogenetic accuracy across layers

We next examined whether phylogenetic accuracy was distributed uniformly across pFM layers. For each protein and network layer, we constructed a consensus phylogeny from the constituent attention-head trees and measured its percentage of correct clades recovered. A distinct pattern emerged for *RAM* phylogenies. The first esm-pFM layer consistently produced the most accurate layer-level phylogenies. Across proteins, first-layer *RAM* consensus phylogenies recovered about 50% of the reference clades on average, substantially more than phylogenies from most later layers (**Fig. 5**). Phylogenetic accuracy declined significantly with increasing layer depth (*p* < 0.02). Later layers continued to produce *RAM*s with substantial phylogenetic signal and contributed correct relationships that the first layer did not always recover. Nevertheless, the high phylogenetic accuracy concentrated in the earliest layer indicates that substantial information useful for distinguishing evolutionary relationships among orthologous proteins is already present in *RAM*s produced by the first transformer layer. But, *SAM* phylogenies showed a different pattern. Their highest accuracies occurred in several intermediate msat-pFM layers, particularly layers 4, 5, 7, and 8 (**Fig. 5**), consistent with previous observations that intermediate msat-pFM layers can effectively extract estimates related to sequence divergence (Lupo et al. 2022).

**Figure 5.**
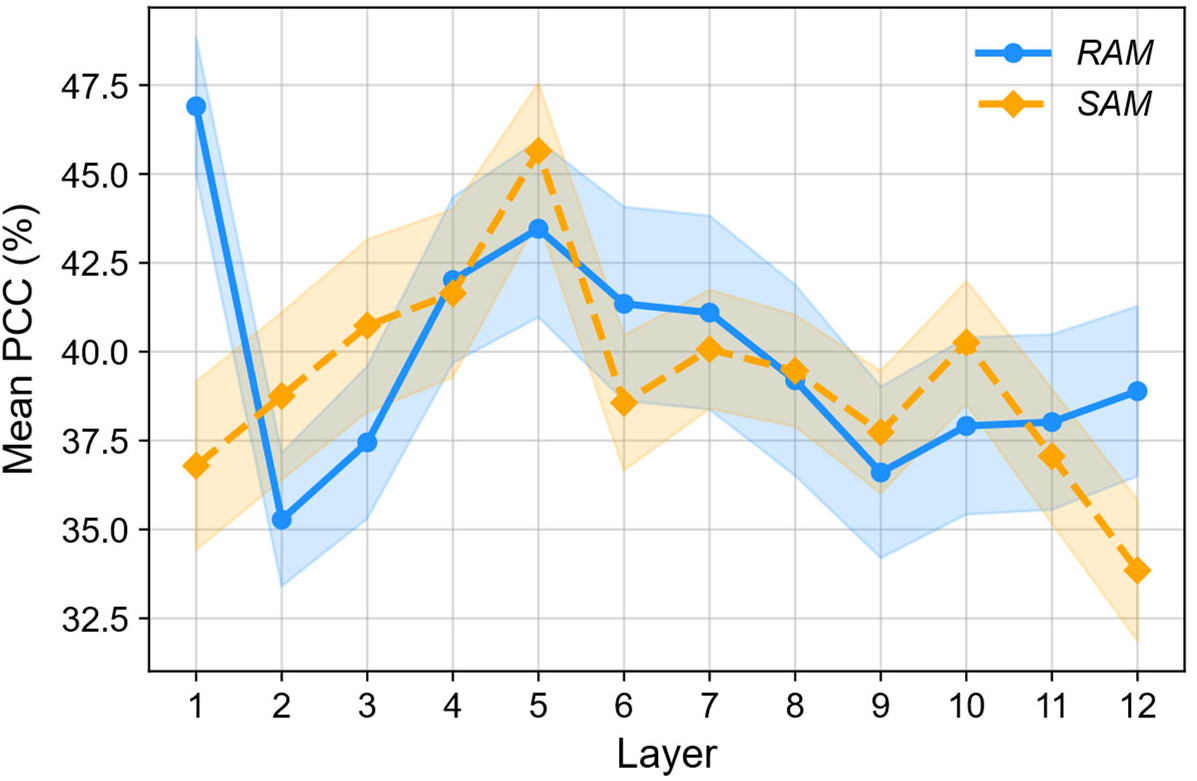
Distribution of phylogenetic information in pFM layers. Mean phylogenetic accuracy (PCC) of layer-consensus phylogenies across the 20 proteins, which are plotted as a function of the esm-pFM (blue) and msat-pFM (orange) layers. The shaded areas indicate the minimum and maximum PCC values across the heads of each layer. Both models had 12 layers (see *Materials and Methods*).

### Comparison of attention-based phylogenies with alternatives

We next compared pFM phylogenies with those inferred using a conventional maximum likelihood (ML) approach. For each of the same 20 proteins, we inferred an ML phylogeny from the MSA using the best-fit amino acid substitution model selected with IQ-TREE 2 (Minh et al. 2020). ML phylogenies were generally more accurate than both *RAM* and *SAM* phylogenies overall (**Fig. 6a**). There is a statistically significant correlation between PCCs of *SAM* phylogenies and the total branch length, indicating that for *SAM*s, faster-evolving proteins tend to result in more accurate trees (correlation = 0.475, p = 0.03). No such trend was observed in *RAM*s (correlation = 0.303, p = 0.19) nor ML (correlation = 0.075, p = 0.75) (**Fig. 6b**). The PCC percentage-point differences between *RAM/SAM* and ML phylogenies also showed a trend: for the 5 slowest evolving proteins, ML had a ΔPCC of +28.8 compared to *RAM*s and ΔPCC of +27.9 compared to *SAM*s. For the 5 fastest evolving proteins, the ΔPCC was only +19.0 and +19.4 respectively. Thus, the performance gap between pFM-based and ML phylogenies was greatest for slowly evolving proteins and narrowed as proteins accumulated more evolutionary change. This pattern suggests that current *RAM* and *SAM* distance procedures require greater sequence divergence to resolve relationships reliably, whereas ML extracts more phylogenetic information from slowly evolving proteins.

**Figure 6.**
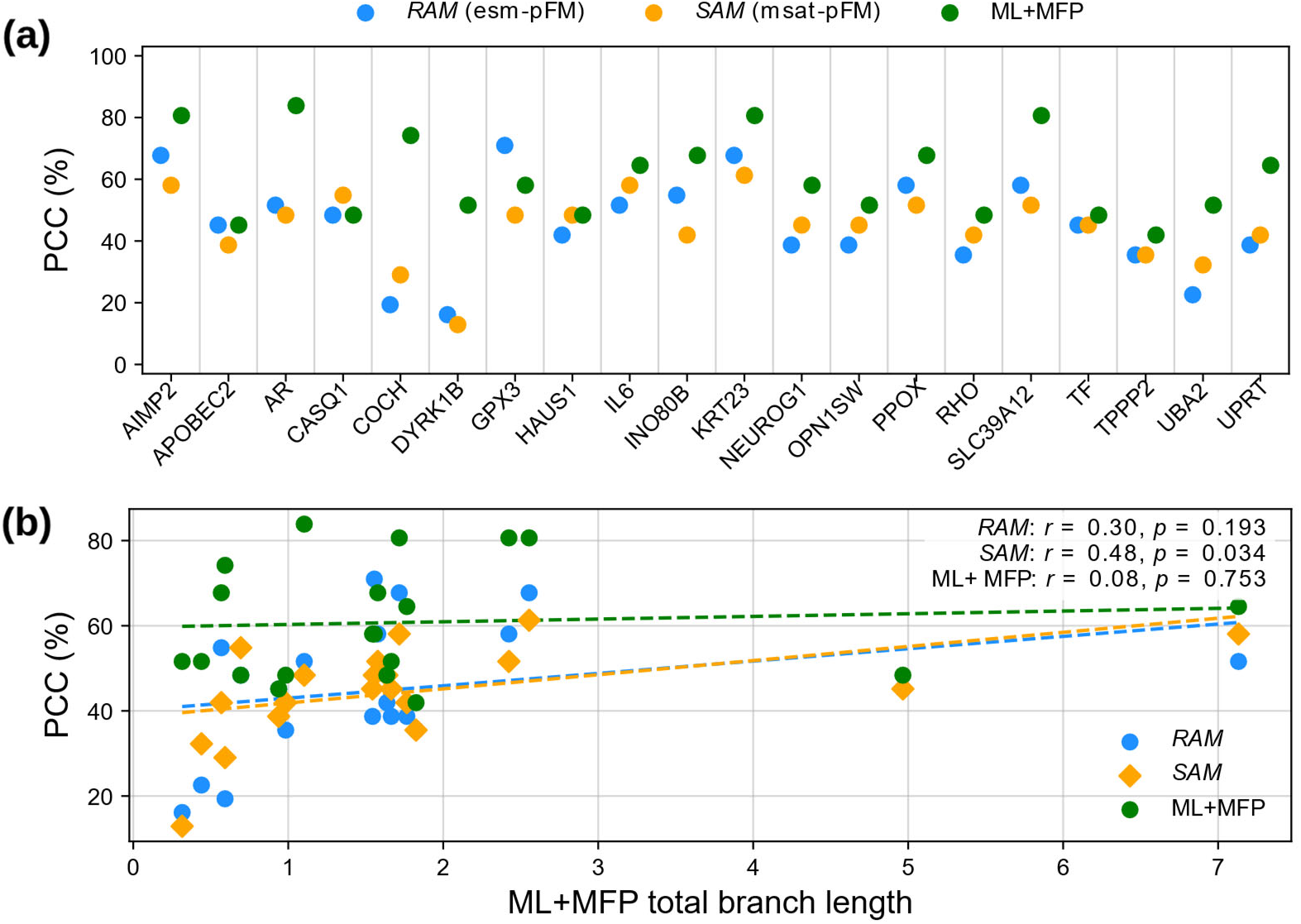
A comparison of *RAM, SAM*, and maximum likelihood (ML) phylogenies. **(a)** Percentage of correct clades recovered in *RAM*, *SAM*, and ML phylogenies for each of the 20 proteins in the 34-species dataset. *RAM* and *SAM* phylogenies were the result of combining all layer/head phylogenies using ASTRAL, whereas the ML phylogeny was obtained using protein-specific best-fit amino acid substitution models. **(b)** Comparison of PCC vs. total branch length as determined by ML, with the Pearson correlation coefficient calculated for each; we find a statistically significant positive correlation (*r*) in *SAM*s, but not in *RAM*s or ML (p-values are shown).

## Discussion

Conventional methods measure differences in amino acids at homologous positions and use amino acid substitution models to infer evolutionary history. *RAM* analysis, in contrast, compares residue dependencies inferred individually for each sequence using a pre-trained esm-pFM. The success of *RAM* phylogenies suggests that residue-dependence patterns also evolve over time, with more closely related species tending to produce more similar *RAM*s. Therefore, classical and pFM-based methods measure different consequences of sequence evolution: conventional evolutionary distances quantify differences in residue states, whereas *RAM* distances quantify accompanying changes in residue context dependence. The ability of *RAM* distances to recover many of the same species relationships as conventional methods suggests that both residue-state differences and changes in residue dependence preserve useful records of the same evolutionary history.

A notable result is that *RAM* phylogenetic accuracy was highest in the first esm-pFM layer and generally declined in subsequent layers. Thus, residue-dependence patterns useful for species phylogeny are already apparent after the earliest stage of sequence analysis. Later layers continue to produce phylogenetically informative *RAM*s, but their average accuracy is lower. In contrast, *SAM* accuracy was greatest in several intermediate msat-pFM layers, consistent with previous observations for sequence-divergence information (Lupo et al. 2022). The different layer profiles further suggest that *RAM*-and *SAM*-based phylogenetic information arises differently during analysis by the two pre-trained pFMs.

Interestingly, however, *SAM* phylogenies did not achieve higher accuracy than *RAM* phylogenies, even though msat-pFM analyzes an explicit MSA and can leverage both sequence and residue dependencies. This contrasts with msat-pFM’s superior performance on other protein structure prediction tasks, particularly secondary structure prediction (Hu et al. 2022). Evoformer, another pFM trained using multiple sequence alignments, also performed better (Hu et al. 2022). These observations suggest that training and inference using MSAs could improve performance by directly incorporating homology information. However, this possibility was not realized in phylogeny inference.

One possible explanation is a mismatch between the evolutionary variation in the alignments used to train the model and the species-phylogeny problem examined here. MSA Transformer was trained using large protein-family alignments assembled from sequence databases. Such alignments reflect variation arising not only from species divergence but also from gene duplication, functional diversification, changes in domain architecture, and deep homology (Rao et al. 2021). Our analyses instead focused specifically on orthologous proteins. *SAM*s summarize relationships among sequences at homologous positions in an alignment, and these relationships may therefore emphasize patterns useful across heterogeneous protein-family alignments during training, but which are not optimal for reconstructing species trees. In contrast, esm-pFM analyzes each ortholog independently, and *RAM* comparison addresses a narrower question: how much has the context-dependent organization of the same protein changed between species?

The pFM approaches are conceptually related to statistical models of residue dependence, such as Potts models (Kumar et al. 2026; Schmelkin et al. 2026). These models estimate pairwise dependencies among residues within a single protein family, thereby allowing the state at one site to depend on the states at other sites. Their main limitation for general phylogenetic analysis is that a separate Potts model must be estimated for each sufficiently large family alignment, and no established methods currently incorporate such models directly into conventional ML phylogenetic analysis. Pre-trained pFMs offer an alternative: a single model with many previously estimated parameters can be applied directly to any protein sequence.

pFMs can be considered to provide a general framework for analyzing context-dependent sequence patterns across protein families, whereas conventional amino acid substitution matrices, such as the Dayhoff family of models, provide general residue-substitution models that typically treat sites independently (Kumar et al. 2026). Whether *RAM* distances can ultimately be given an explicit evolutionary interpretation comparable to substitutions per site remains unresolved. Nevertheless, their phylogenetic performance suggests that context-dependent sequence information can be harnessed without estimating a new family-specific dependency model for every protein, which may not be feasible.

*RAM* phylogenies do not explicitly assume a stationary or reversible substitution process, nor do they assume independent site evolution, as is common in traditional statistical phylogenetics. This makes them potentially attractive for proteins in which residue context, compositional shifts, or lineage-specific constraints violate conventional model assumptions, and investigating such cases will be an important direction for future work. Nevertheless, the absence of an explicit substitution model cannot be equated with the absence of assumptions. Protein foundation models are influenced by their training-data composition, sequence redundancy, taxonomic sampling, and other training-corpus properties (Bernett et al. 2024; Li et al. 2024). These assumptions are implicit rather than specified as an evolutionary rate matrix, but they can still affect the phylogenetic inferences these pFMs make.

Foundation model phylogenetics also currently lacks several features that established statistical approaches provide. *RAM* distances do not yet yield interpretable estimates of the number of substitutions, and no likelihood-based framework exists for comparing alternative phylogenetic hypotheses. Branch lengths inferred from attention distances therefore cannot yet be interpreted as substitutions per site. Statistical procedures for assessing confidence in *RAM* phylogenies also remain to be developed. Attention-based approaches are therefore best viewed as complementary to conventional phylogenetic methods at present, particularly for analyses in which individual homologous sequences can be examined directly without constructing a multiple sequence alignment (Kumar et al. 2026).

## Conclusions

Analyses of orthologous proteins using pre-trained foundation models reveal that residue-attention patterns contain substantial information about species relationships. *RAM* distances obtained after individual sequences were analyzed with esm-pFM were as accurate as *SAM* distances obtained after MSAs were analyzed with msat-pFM. This equivalence persisted across distantly related species, closely related great apes, and larger mammalian datasets. Much of the *RAM* phylogenetic signal was concentrated in the earliest esm-pFM layer, and some of the *RAM* phylogenies approached the accuracy of conventional maximum likelihood analyses for the proteins examined.

These findings suggest that protein evolution leaves a measurable record not only in differences among homologous amino acids but also in changes in the context-dependent relationships among residues. *RAM* phylogenetics currently lacks interpretable evolutionary branch lengths, statistical confidence procedures, and an explicit model of sequence evolution, so it is not a replacement for conventional phylogenetic inference. Nevertheless, it offers a fundamentally different route to infer species relationships from proteins and establishes residue attention as a useful new quantity for molecular evolutionary analysis.

## Materials and Methods

### OrthoMaM data

We obtained a well-curated set of protein sequences from the Orthologous Mammalian Markers (OrthoMaM v12) database (Allio et al. 2024). We sampled a set of proteins shorter than 1024 residues, a limit imposed by msat-pFM. From this set, we selected 20 proteins containing representative sequences for three species sets: a set of 4 distantly related mammals (*Homo sapiens, Macaca fascicularis, Rattus norvegicus, Monodelphis domestica*), a set of 4 closely related great ape species (*Homo sapiens, Pan troglodytes, Gorilla gorilla,* and *Pongo abelii*), and a larger set of 34 species representing 10 mammalian orders (see *Supplementary Table S1*). We chose the 34 species to ensure each had a representative sequence for each of the 20 selected proteins, and they included the same species from the 4-mammals and 4-apes sets.

We also used the OrthoMaM SuperMatrix tree, pruned to the 34 species of interest, as the reference tree. Because the OrthoMaM SuperMatrix tree was assembled from a much larger pool of >15K CDS markers, we consider reconstructing it from only 20 proteins an adequate proxy for the accuracy of *RAM* and *SAM* phylogenetic inference methods.

### pFM models

We downloaded the pretrained esm-pFM and msat-pFM models from the fair-esm GitHub repository and saved them as PyTorch objects for independent downstream analysis. We chose the 35M-parameter esm-pFM checkpoint to obtain a model smaller than the 100M-parameter msat-pFM (128 Mb vs. 442 Mb), to rule out the effect of a higher parameter count on esm-pFM’s competitive performance. However, larger parameter models did not significantly improve phylogeny accuracy with esm-pFM (results not shown).

### Phylogeny inference from *RAM*s

For the esm-pFM analyses, we removed the gap characters (“-”) from every sequence in each alignment, resulting in FASTA files with unaligned sequences. We used these unaligned sequences to infer *RAM*s individually and then reconstructed phylogenetic trees using the *RAM*-based method described in the Results section. Each protein analysis produced 240 *RAM* trees, as the 35M-parameter esm-pFM checkpoint we used has 12 attention layers, each with 20 attention heads. The computational runtime for this process across different protein lengths is listed in Supplementary Figure S1.

### Phylogeny inference from *SAM*s

For the msat-pFM analyses, we used msat-pFM to generate *SAM*s directly from MSAs. We inferred phylogenetic trees from these *SAM*s for each of the three protein sets (4 mammals, 4 apes, and 34 mammalian species). The msat-pFM model used here contains 12 attention layers with 12 attention heads per layer, yielding 144 layer/head combinations for each input MSA. Unlike *RAM* inference, which analyzes each protein sequence independently and thus requires pairwise sequence alignment as an intermediary step, *SAM* inference uses the MSA directly (**Fig. 1b**).

For an MSA containing *N* species and *L* alignment columns, each layer/head of msat-pFM produces an *N* × *N* sequence attention matrix (*SAM*) at every alignment column. Each row of a *SAM* describes the distribution of attention from one species sequence to all species sequences in the MSA at that site. For each species pair A and B, we calculated the Hellinger distance between their corresponding rows separately at every alignment column. We then combined these column-specific Hellinger distances by calculating their root-mean-square (RMS) across all *L* alignment columns, yielding a single pairwise distance *d*_AB_ for that layer/head. Repeating this procedure for all species pairs produced one symmetric *N* × *N* pairwise distance matrix for each layer/head. We used the lower triangle of each matrix to infer an NJ phylogeny, yielding 144 phylogenies per protein. We combined the resulting layer/head phylogenies using ASTRAL to infer the protein-level consensus phylogeny. Because no repeated pairwise alignment step is required, the computational runtime for *SAM* tree-building was much faster than for *RAM* tree-building (see *Supplementary Figure S1*).

### Inferring maximum likelihood phylogenies for comparison

To benchmark classical maximum-likelihood (ML) analysis, we used IQ-TREE 2 (Minh et al. 2020). Using the same 20 protein MSAs from OrthoMaM, we inferred an ML phylogeny for each protein using the best-fit amino acid substitution model selected by ModelFinder Plus. We again used ASTRAL to derive a 20-protein phylogeny for the ML+MFP trees.

### Assessing the accuracy of phylogenetic inference for larger trees

We quantified each phylogeny’s accuracy using the Robinson-Foulds (Robinson and Foulds 1981) distance from the OrthoMaM reference tree, derived from the entire OrthoMaM database (rather than just the 20 proteins in our sample), as a proxy for a ground-truth tree (Allio et al. 2024). We used this to calculate a percentage metric, PCC, which is the percentage of clades inferred correctly, as determined by the equation:

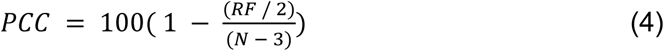

where *RF* is the Robinson-Foulds distance, and *N* is the number of species in the tree.

## Supplementary Material

Supplementary material will be available online at *Molecular Biology and Evolution*.

## Acknowledgments

The authors thank Dr. Sudip Sharma for his advice and technical assistance.

## Data Availability

The supplemental information, datasets, and scripts are available at https://github.com/RohanAlibutud/PLM-Phylo.git. The foundation models referenced are available at https://github.com/facebookresearch/esm.

## Supplementary Information

**Supplemental Table S1-.** 34 mammalian species used in this analysis.

| Scientific Name | Common Name |
| --- | --- |
| *Acomys russatus* | Golden spiny mouse |
| *Bos indicus* | Zebu |
| *Bos taurus* | Domestic cattle |
| *Camelus dromedarius* | Dromedary camel |
| *Camelus ferus* | Wild Bactrian camel |
| *Cercocebus atys* | Sooty mangabey |
| *Equus caballus* | Horse |
| *Felis catus* | Domestic cat |
| *Gorilla gorilla gorilla* | Western lowland gorilla |
| *Heterocephalus glaber* | Naked mole-rat |
| *Homo sapiens* | Human |
| *Ictidomys tridecemlineatus* | Thirteen-lined ground squirrel |
| *Macaca fascicularis* | Crab-eating macaque |
| *Manis javanica* | Sunda pangolin |
| *Marmota flaviventris* | Yellow-bellied marmot |
| *Marmota marmota marmota* | Alpine marmot |
| *Mesocricetus auratus* | Golden hamster |
| *Microtus ochrogaster* | Prairie vole |
| *Molossus molossus* | Velvety free-tailed bat |
| *Monodelphis domestica* | Gray short-tailed opossum |
| *Mus musculus* | House mouse |
| *Myotis brandtii* | Brandt's bat |
| *Myotis lucifugus* | Little brown bat |
| *Myotis myotis* | Greater mouse-eared bat |
| *Ochotona princeps* | American pika |
| *Ovis aries* | Domestic sheep |
| *Pan troglodytes* | Chimpanzee |
| *Peromyscus californicus insignis* | California mouse |
| *Pongo abelii* | Sumatran orangutan |
| *Rattus norvegicus* | Norway rat |
| *Saimiri boliviensis boliviensis* | Bolivian squirrel monkey |
| *Sus scrofa* | Wild boar / Domestic pig |
| *Tupaia chinensis* | Chinese tree shrew |
| *Vulpes vulpes* | Red fox |

**Supplementary Figure S1:**
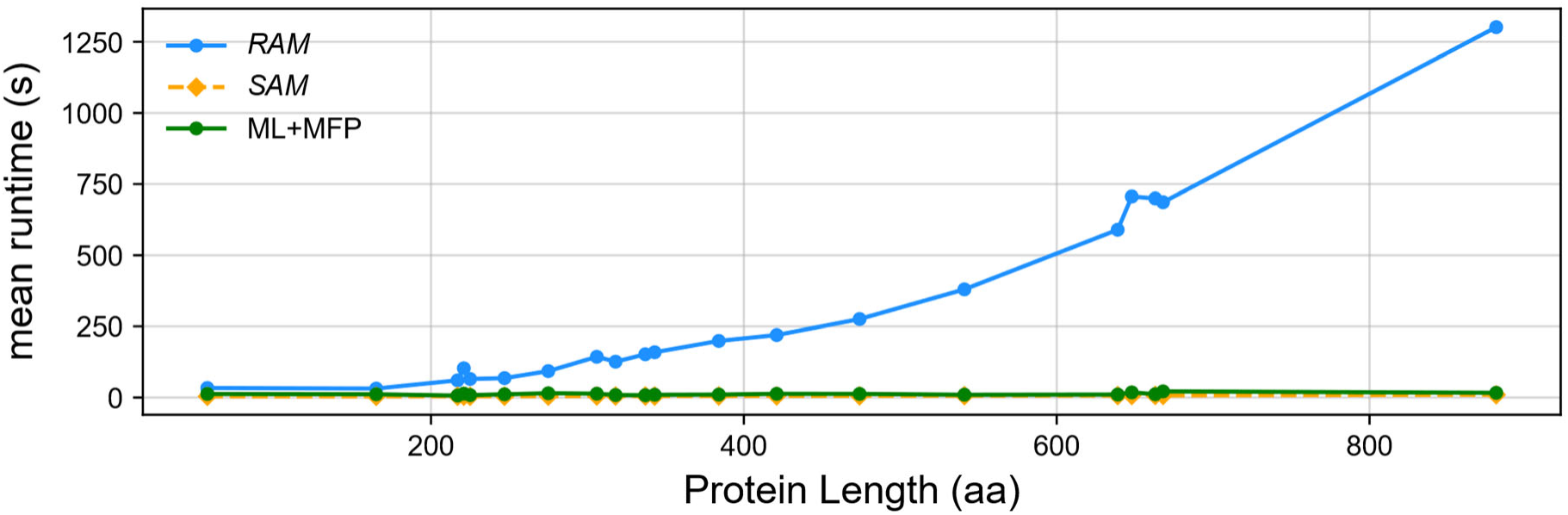
Computational runtime for pFM tree-building. Computational time required for single-threaded *RAM*, *SAM*, and ML analysis on a personal computer with 64 GB of RAM and 32 processors.

